# Sensorimotor characteristics of sign translations modulate EEG when deaf signers read English

**DOI:** 10.1101/337170

**Authors:** Lorna C. Quandt, Emily Kubicek

## Abstract

Bilingual individuals automatically translate written words from one language to another. While this process is established in spoken-language bilinguals, there is less known about its occurrence in deaf bilinguals who know signed and spoken languages. Since sign language uses motion and space to convey linguistic content, it is possible that action simulation in the brain’s sensorimotor system plays a role in this process. We recorded EEG from deaf participants fluent in ASL as they read individual English words and found significant differences in alpha and beta EEG at central electrode sites during the reading of English words whose ASL translations use two hands, compared to English words whose ASL translations use one hand. Hearing non-signers did not show any differences between conditions. These results demonstrate the involvement of the sensorimotor system in cross-linguistic, cross-modal translation, and suggest that action simulation processes may be key to deaf signers’ language concepts.

## 1. Background

Cross-linguistic translation occurs when a bilingual person automatically accesses mental lexicons for both known languages even when translation is not required (Baten, Hofman, & Loeys, 2010; Duyck, Van Assche, Drieghe, & Hartsuiker, 2007; Lemhöfer & Dijkstra, 2004). Substantial evidence demonstrates cross-linguistic translation in spoken languages among bilinguals who use two spoken languages (unimodal bilinguals; Midgley, Holcomb, & Grainger, 2009; Thierry & Wu, 2007). Researchers have made comparatively less progress regarding how deaf bilinguals link meaning between languages that rely on different modalities—for example, written language (e.g., English) and sign language (e.g., American Sign Language, ASL). It is possible that while processing written English, deaf bilinguals activate the corresponding ASL translations. Both behavioral and psychophysiological evidence suggests that automatic cross-linguistic translation does occur when deaf signers read English (Giezen, Blumenfeld, Shook, Marian, & Emmorey, 2015; Giezen & Emmorey, 2015; Meade, Midgley, Sehyr, Holcomb, & Emmorey, 2017; Morford, Wilkinson, Villwock, Piñar, & Kroll, 2011). These researchers use phonological interference paradigms, presenting pairs of written words, some of which have ASL translations that share phonological properties (e.g., location of sign). Thus, participants in these studies may become aware of the experimental manipulation, possibly using conscious strategies to complete the experimental task (Meade et al., 2017). The canonical language networks in the brain have been identified as a likely mechanism for how this phenomenon occurs (Meade et al., 2017).

Since sign language uses motion and space to convey linguistic content, an individual who is fluent in sign language may implicitly draw upon their brain’s sensorimotor system during cross-linguistic, cross-modal translation. A consideration of ASL-related processing without discussion of the sensorimotor system of the brain is incomplete (Corina, Lawyer, & Cates, 2013; Emmorey, McCullough, Mehta, & Grabowski, 2014; Gutierrez-Sigut, Payne, & MacSweeney, 2016). For instance, while perceiving ASL is likely to rely on some modality-independent processing, such as the recruitment of classical phonology regions during ASL phonology tasks (Petitto et al., 2016), recent work shows that hearing interpreters process sign language phonology by activating visuo-motor systems to relay gestural input to the language network (Kanazawa et al., 2017). Since ASL signs are produced with the hands and arms, producing ASL is associated with increased activity in the primary sensory and motor cortices (Emmorey et al., 2014; Emmorey, Mehta, McCullough, & Grabowski, 2016).

Thus, we know that the sensorimotor systems are involved in sign processing to some extent. But, we don’t know how the sensorimotor system is involved in cross-modal, cross-linguistic translation in deaf readers. Embodied cognition theorists posit that perception is grounded in one’s own ability to act (Barsalou, 2008; Creem-Regehr & Kunz, 2010; Gallese & Sinigaglia, 2011). One instantiation of this theory is the action observation system (AON) of the human brain, which includes brain regions that are uniquely sensitive to other people’s actions (Caspers, Zilles, Laird, & Eickhoff, 2010; Molenberghs, Cunnington, & Mattingley, 2012). The AON is thought to include the putative human mirror neuron system, which has been a topic of much debate (Gallese, Gernsbacher, Heyes, Hickok, & Iacoboni, 2011; Hickok, 2009; Kessler & Garrod, 2013; Rizzolatti & Sinigaglia, 2010, 2016). One likely function of the mirroring system, and of the AON, is action simulation—a phenomenon wherein somatosensory and motor aspects of an action are simulated even in the absence of action production. For instance, an observer of dance may draw upon her own sensorimotor system to simulate the movements involved in another person’s dance routine (Calvo-Merino, Glaser, Grezes, Passingham, & Haggard, 2005; Calvo-Merino, Grèzes, Glaser, Passingham, & Haggard, 2006; Gardner, Goulden, & Cross, 2015; Kirsch & Cross, 2015), or a person may mentally simulate an imagined action (Decety, 1996; Jiang, Edwards, Mullins, & Callow, 2015).

Action simulation results in changes in the sensorimotor alpha (8-13 Hz) and beta (~14-25 Hz) rhythms present within the EEG signal. Typically, imagining or observing actions leads to a suppression, or desynchronization, of alpha and beta rhythms measurable at electrodes over the primary somatosensory and primary motor cortices (Avanzini et al., 2012; de Lange, Jensen, Bauer, & Toni, 2008; Muthukumaraswamy, Johnson, & McNair, 2004; Quandt & Marshall, 2014). When measured at central electrodes overlying the central sulcus, 8-13 Hz alpha suppression is most linked to activity within the primary somatosensory cortex, and 14-22 Hz beta suppression is more correlated with activation of the primary motor cortex (Ritter, Moosmann, & Villringer, 2009; Salmelin & Hari, 1994).

A growing body of literature has examined the functionality of the AON and mirroring-related regions in sign language processing, given the visual-spatial movements required to produce signs, and the action observation processes which are likely to be involved in sign perception. Using functional magnetic resonance imaging (fMRI), a number of studies have shown little to no involvement of classical “mirroring” processes during deaf signers’ perception of ASL (Corina & Gutierrez, 2016; Corina & Knapp, 2006; Emmorey, Xu, Gannon, Goldin-Meadow, & Braun, 2010). Emmorey and colleagues (2010) examined the differences in mirroring activity between hearing non-signers and deaf signers during observation of pantomimes (e.g. peeling an imaginary banana) and ASL action verbs. Results showed activations for pantomimed actions and ASL verbs in regions associated with mirror-related processing in hearing non-signers. Deaf signers showed no significant activations in these mirroring areas when viewing pantomimes. These results echo that of Corina et al. (2006), which indicate insignificant recruitment of the AON for deaf signers during observation of intransitive self-oriented actions (e.g. rubbing one’s eyes), transitive object-oriented actions (e.g. opening a door), and ASL signs, while fronto-parietal activations were observed for hearing non-signers under these same conditions. These results suggest that hearing non-signers recruit the AON when observing ASL while deaf signers do not. MacSweeney et al.’s (2004) work on gesture and sign language, however, demonstrates that deaf signers recruit parts of the AON more robustly during sign language perception compared to a non-language gestural communication system. In contrast, hearing non-signers showed the opposite effect (communication system > BSL) in similar AON areas. Taken together, these studies indicate that more research exploring the interaction between bimodal language experience and the AON is needed.

### 1.1 The current study

Drawing upon existing work, we designed a study to assess whether implicit cross-modal, cross-linguistic translation would affect sensorimotor EEG rhythms, indicating the involvement of the sensorimotor system in the translation occurring when deaf signers read English. If the sensorimotor system is involved in this process, we would be able to conclude that action simulation processes occur during reading, wherein deaf readers unconsciously simulate ASL translations of English words.

Functional neuroimaging has shown that the right primary sensory and motor cortices are more active when deaf signers produce two-handed signs, compared to one-handed signs (Emmorey et al., 2016), a pattern which is in line with existing work using non-linguistic movements (Toyokura, Muro, Komiya, & Obara, 2002). If direct action simulation processes were involved during cross-linguistic, cross-modal translation processes, we expect that English words with a two-handed ASL translation would result in greater alpha and beta desynchronization over the right pre- and post-central region.

One line of work has asked whether reading action-related language invokes action simulation processes, in the same way as observing action does. While there is some divergence regarding this hypothesis (Kemmerer & Gonzalez-Castillo, 2010; Postle, McMahon, Ashton, Meredith, & de Zubicaray, 2008), a number of studies have demonstrated that reading action words results in somatotopic activation of sensorimotor cortices, suggesting that implicit simulation of action may contribute to a reader’s comprehension of a written word (Hauk, Johnsrude, & Pulvermüller, 2004; Hauk & Pulvermuller, 2011; Pulvermüller, Härle, & Hummel, 2001; Schaller, Weiss, & Müller, 2017). The current study draws upon this work to examine sensorimotor EEG rhythms during single word reading.

We collected EEG from across the scalp as deaf signers and hearing non-signers read English words for which the ASL translations used either one or two hands. The design allowed us to compare sensorimotor involvement between words with one- and two-handed ASL translations for each group. Our task involved no mention of ASL, and there was no deliberate interference stemming from ASL phonology. This design removed the issue of awareness of ASL phonology during the reading task, allowing for an entirely passive approach. The work we describe here is the first time-frequency EEG analysis of neural activity while deaf signers read.

We predicted that deaf signers would show significant differences in alpha and beta EEG rhythms during the reading of English words whose ASL translations use either 1 or 2 hands. Specifically, we expected that the increased sensorimotor demands of carrying out a 2-handed sign would be reflected in the simulated neural activity, resulting in differences over the right fronto-central regions overlying primary sensorimotor cortices and other key regions of the AON (e.g., premotor cortex and supplementary motor area). The right laterality of this effect was predicted since a key difference between a 1-handed ASL sign (always produced with the dominant hand) and a 2-handed ASL sign is the use of the left hand. We expected that both the alpha rhythm of the EEG signal (~8-13 Hz) and the beta rhythm (~14-25 Hz) would show greater desynchronization in response to 2-handed signs. We were less confident in the direction of the effect for the beta rhythm, given the complex nature of the beta rhythm (Spitzer & Haegens, 2017). We expected that the hearing group would show no significant differences between conditions, given that they do not know ASL and thus there should be no automatic translation processes occurring in this group.

## 2. Materials and Methods

### 2.1 Participants

Twenty-four Deaf fluent signers (12 females; 1 declined to respond) and twenty-two Hearing non-signers (8 females; 1 declined to respond) participated in the experiment (mean age = 28.9 years, SD = 8.99). Participants signed an informed consent form presented in written English and American Sign Language that had been approved by the university Institutional Review Board. All Deaf signers were self-identified as fluent in American Sign Language with 91.6% reporting they started using ASL at or before the age of 5 (mean = 2.88, SD = 5.36, median = 1). The hearing non-signers reported normal hearing and no knowledge of any signed language. All participants self-reported as neurologically healthy, having normal or corrected vision, and all but two reported being right-handed. Educational, demographic, and language background information is shown in Tables 1 and 2.

**Table 1.** Formal education. Self-reported highest educational degree obtained for Deaf and Hearing participants.

|  | Deaf | Hearing |
| --- | --- | --- |
| Some high school | 0 | 0 |
| High school | 4 (16.7%) | 0 |
| GED | 1 (4.2%) | 0 |
| Some college | 3 (12.5%) | 1 (4.5%) |
| Associates | 0 | 2 (9.1%) |
| Bachelors | 4 (16.7%) | 7 (31.8%) |
| Some grad school | 2 (8.3%) | 3 (13.6%) |
| Masters | 6 (25%) | 8 (36.4%) |
| Doctorate | 4 (16.7%) | 1 (4.5%) |

**Table 2.** Demographics and language background information for all participants.

|  | Deaf | Hearing | t-test *p* value |
| --- | --- | --- | --- |
| Age (years) | 30.8 (11.6) | 27.1 (4.5) | .16 |
| Current written English use <sup>a</sup> | 1.5 (.9) | 1.3 (.6) | .41 |
| % Written English understanding <sup>b</sup> | 87.8 (17.5) | 93.9 (21) | .30 |
| % Written English production <sup>b</sup> | 81 (20.7) | 91 (21) | .09 |
| Current ASL use <sup>a</sup> | 1.3 (.5) |  |  |
| ASL understanding <sup>b</sup> | 90.8 (10.9) |  |  |
| ASL production <sup>b</sup> | 95 (6.2) |  |  |

<sup>a</sup> Self-reported rating on a 7-point scale ranging from 1 = "All the time" to 7 = "Only on special occasions (e.g. home for the holidays)".
<sup>b</sup> Self-rated proficiency on a sliding scale from 0 (poorly) – 100 (fluently).

### 2.2 Stimuli

We created two words lists, each containing 40 words. One list consisted of English words for which the ASL translation uses only one hand (‘one-handed’ or 1H words; e.g., warm); the other list contained English words for which the ASL translation uses two hands (‘two-handed’ or 2H words; e.g., family). We selected the words by choosing ASL signs from the ASL-Lex database (Caselli, Sehyr, Cohen-Goldberg, & Emmorey, 2016) whose English translations fulfilled our criteria. None of the words selected were verbs; the stimulus set included nouns, adjectives, and adverbs. We aimed to select words which had a very reliable ASL translation, avoiding words with many variations of ASL translations. This was determined in consultation with deaf experimenters fluent in ASL.

There were no significant differences between the English words contained in 1H and 2H lists for any of the following measures: word length, frequency, number of phonemes, imageability, mean reaction time and standard deviation during a lexical decision task, and mean reaction time and standard deviation during a naming task (Balota et al., 2007; MRC Psycholinguistic Database: Machine Usable Dictionary. Version 2.00; Kučera & Francis, 1967). See Table 3 for summary statistics.

There were no significant differences between the ASL sign translations of the English words contained in the 1H and 2H lists for any of the following measures of the signs: frequency in ASL, iconicity, and flexion (Caselli et al., 2016). See Table 3 for summary statistics.

An additional 12 animal words were included in the experiment, for use as “catch” trials. The data related to these animal words was not analyzed at any point.

**Table 3.** English and ASL norms for 2 categories of stimuli: 1-handed (1H) words, and 2-handed (2H) words.

|  |  | 1H words: M (SD) | 2H words: M (SD) | t-test *p* value |
| --- | --- | --- | --- | --- |
| English norms | Word length | 5.20 (1.67) | 5.72 (2.17) | .23 |
|  | Frequency (K&F) | 102.36 (151.44) | 108.93 (142.30) | .84 |
|  | Log frequency (HAL) | 9.62 (1.69) | 10.01 (1.44) | .27 |
|  | # phonemes | 4.09 (1.11) | 4.55 (1.34) | .10 |
|  | Lexical Decision RT M | 605.55 (45.50) | 617.54 (62.97) | .33 |
|  | Lexical Decision RT SD | 202.14 (63.97) | 217.57 (87.57) | .37 |
|  | Naming RT M | 605.47 (42.57) | 607.26 (42.74) | .85 |
|  | Naming RT SD | 127.97 (58.10) | 127.89 (52.68) | .99 |
|  | Imageability | 509.47 (112.49) | 541.97 (80.48) | .20 |
| ASL norms | Frequency (ASL-Lex) | 4.74 (.75) | 4.49 (1.01) | .22 |
|  | Iconicity | 3.71 (1.68) | 3.73 (1.34) | .97 |
|  | Flexion | 3.38 (2.20) | 3.63 (2.18) | .61 |

### 2.3 Experimental Design

Stimuli were individual English words presented in the center of 27” computer monitor approximately 2.5 feet from the participant. Word stimuli were presented in four blocks with a break after each block. Each block included 23 words consisting of 1-handed, 2-handed, and animal words that were pseudorandomized. Each word was presented once during the experiment, for a total of 40 1H trials and 40 2H trials. Participants were instructed to count how many animal words they observed in one block and reported their answer to the experimenter during the breaks between blocks. These responses were not recorded, as the purpose of this task was to ensure participant alertness throughout the study. If a participant gave any significantly deviant responses, experimenters would check in with participants to ensure they were attending to the experiment. This did not occur for any participants.

### 2.4 Recording

EEG was recorded from 64 active Ag/AgCl electrodes using an actiCAP setup (Brain Products GmbH, Germany), in combination with SuperVisc electrode gel. Data were recorded using an online Cz reference and an AFz ground. Saline gel (SuperVisc, Brain Products) was inserted into each electrode to lower impedances below 25 kΩ. The EEG signals were amplified by the individual electrode amplifiers, and again by a 24-bit actiCHAMP amplifier (Brain Vision LLC, Morrisville, NC). Hardware filter settings included a high-pass filter (.53 Hz) and a low-pass filter (120 Hz). Data was collected at a 1000 Hz sampling rate.

### 2.5 Data Preparation

All data processing was implemented using EEGLAB v. 14.1.1 (Delorme & Makeig, 2004). Data were referenced offline to the average of the two mastoid electrodes (TP9 and TP 10). Data were filtered offline using a .1 Hz high-pass and a 100 Hz low-pass filter. Epochs were extracted from the continuous EEG, time locked to each stimulus of interest (all 1-handed, 1H, and 2-handed words, 2H). Onset of the stimulus was considered time 0, and epochs included data from −1.5 s to 2.5 surrounding each stimulus onset. A baseline period was defined as −200 to 0 s prior to stimulus onset. We ran an independent component analysis (ICA) on each epoched dataset (two datasets per participants: 1H and 2H). A trained experimenter visually inspected ICA results for each dataset and removed ICs deemed to be eyeblink artifacts or other non-brain artifacts (mean and mode number of ICs removed per participant = 1). Trials were removed from further analysis if they included large amounts of artefactual activity, as judged by a highly-trained experimenter (total # of trials removed: 3 1H for one participant; 3 2H for one participant and 1 2H for 3 participants). All datasets were compiled into one study folder and assigned groups (Deaf and Hearing) and conditions (1H and 2H).

### 2.6 Time-Frequency Analysis

Event-related spectral perturbation (ERSP) was computed at each electrode within our central region of interest (ROI), conducting a paired t-test (1H vs. 2H) for each group (Deaf and Hearing).

We conducted planned t-tests to obtain a full picture of the characteristics of the data, driven by a priori predictions developed before data analysis. For these planned tests, we analyzed all scalp electrodes at four frequency bands: low alpha (8-10 Hz), high alpha (11-13 Hz), low beta (14-17 Hz) and high beta (18-25 Hz). For each frequency band, we divided the epoch of interest into four bins, encompassing the entire time the stimulus was present on the screen: 0-250 ms, 250-500 ms, 500-750 ms, and 750-1000 ms. For each analysis, we considered an effect to be of interest if it was significant at three adjacent electrodes. To control for multiple comparisons, we used a Bonferroni corrected *p* values of .016 (.05 / 3) as the threshold for significance.

To get a more fine-grained view of the time and frequency dynamics of sensorimotor EEG during word perception, we calculated ERSPs at 21 electrodes comprising a region of interest over the pre- and post-central gyrus. The 21 electrodes included were FC_5_, FC_3_, FC_1_, FCz, FC_2_, FC_4_, FC_6_, C_5_, C_3_, C_1_, Cz, C_2_, C_4_, C_6_, CP_5_, CP_3_, CP_1_, CPz, CP_2_, CP_4_, and CP_6_. At each of these electrodes we compared time-frequency plots (from 0-1000 ms in time and 8 to 25 Hz in frequency, encompassing the alpha and beta ranges) between 1-handed and 2-handed words for both the Deaf and Hearing groups. We were particularly interested in significant effects seen at these electrodes since alpha and beta rhythms present at central electrodes are closely tied to activity in pre- and post-central gyri (primary motor and primary somatosensory cortices, respectively), which are key components of the AON (Arnstein, Cui, Keysers, Maurits, & Gazzola, 2011; Perry & Bentin, 2009; Ritter et al., 2009). For these region-of-interest analyses we used a *p* value of .05, with a false-discovery rate correction applied to control for false positives.

## 3. Results

### 3.1 Alpha and beta activity across the scalp

No significant differences between 1H and 2H words were observed at any time bin in the lower alpha band (8-10 Hz) for either Deaf or Hearing groups.

In the upper alpha band (11-13 Hz), there was a significant interaction effect (*p* < .016) between the Groups and the Conditions. The significant interaction effect was observed at 5 frontal electrodes (F_7_, AF_7_, FP_1_, FP_2_, and AF8) from 500-750 ms following stimulus onset. No pairwise comparisons yielded significant results, so this effect was not further interpreted. No significant differences were observed at any other time bin.

In the lower beta band (14-17 Hz), there was a significant Group × Condition interaction in all four time bins spanning 0-1000 ms post stimulus onset. In the 250-500, 500-750, and 750-1000 ms bins, pairwise comparisons revealed significant differences in low beta band power between Deaf and Hearing groups, for observation of 2-handed words only. In all three time bins, the significant effects were observed at left frontal electrodes and reflected higher beta power in the Deaf group when reading 2-handed words, compared to the Hearing group. No significant differences were seen between conditions for the Hearing group.

In the upper beta band (18-25 Hz), there was a significant Group × Condition interaction observed during each of the four time bins spanning 0-1000 ms post stimulus onset. For all four time bins, pairwise comparisons revealed a significant difference in high beta power during observation of one-handed vs. two-handed words, for the Deaf group only. In all time bins, the significant effects were observed at fronto-central electrodes (see Figure 2), reflecting higher beta power during the reading of two-handed words compared to one-handed words. In the 500-750 ms bin, a significant difference was also observed over right parieto-occipital sites, driven by higher beta power in this area for 2-handed words. No significant differences were seen between conditions for the Hearing group.

**Figure 1.**
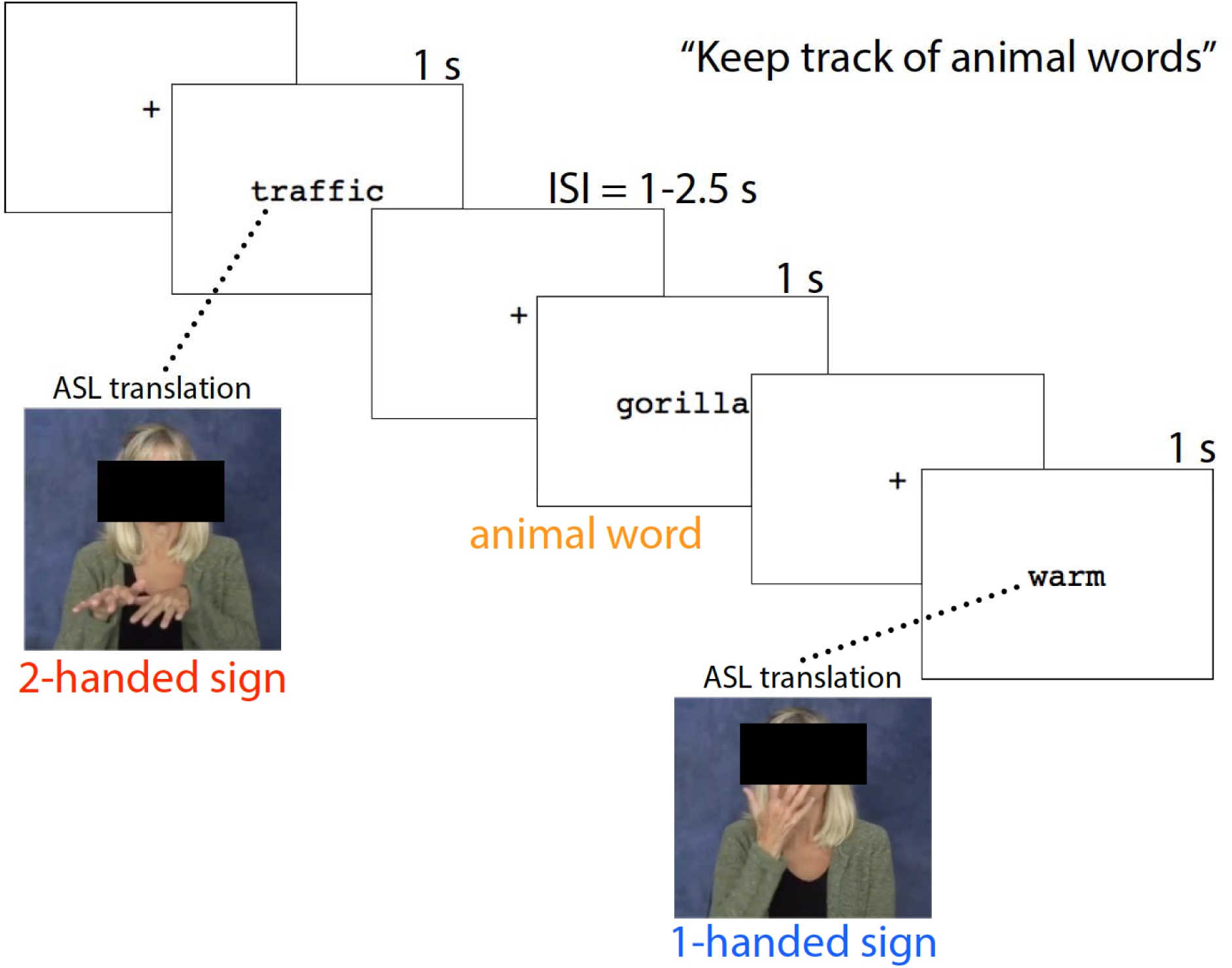
Overview of experimental task. Participants saw one word at a time displayed on a screen and were instructed to mentally count animal words (catch trials). Trials of interest were English words whose ASL translation required either one hand (1H) or two hands (2H).

**Figure 2.**
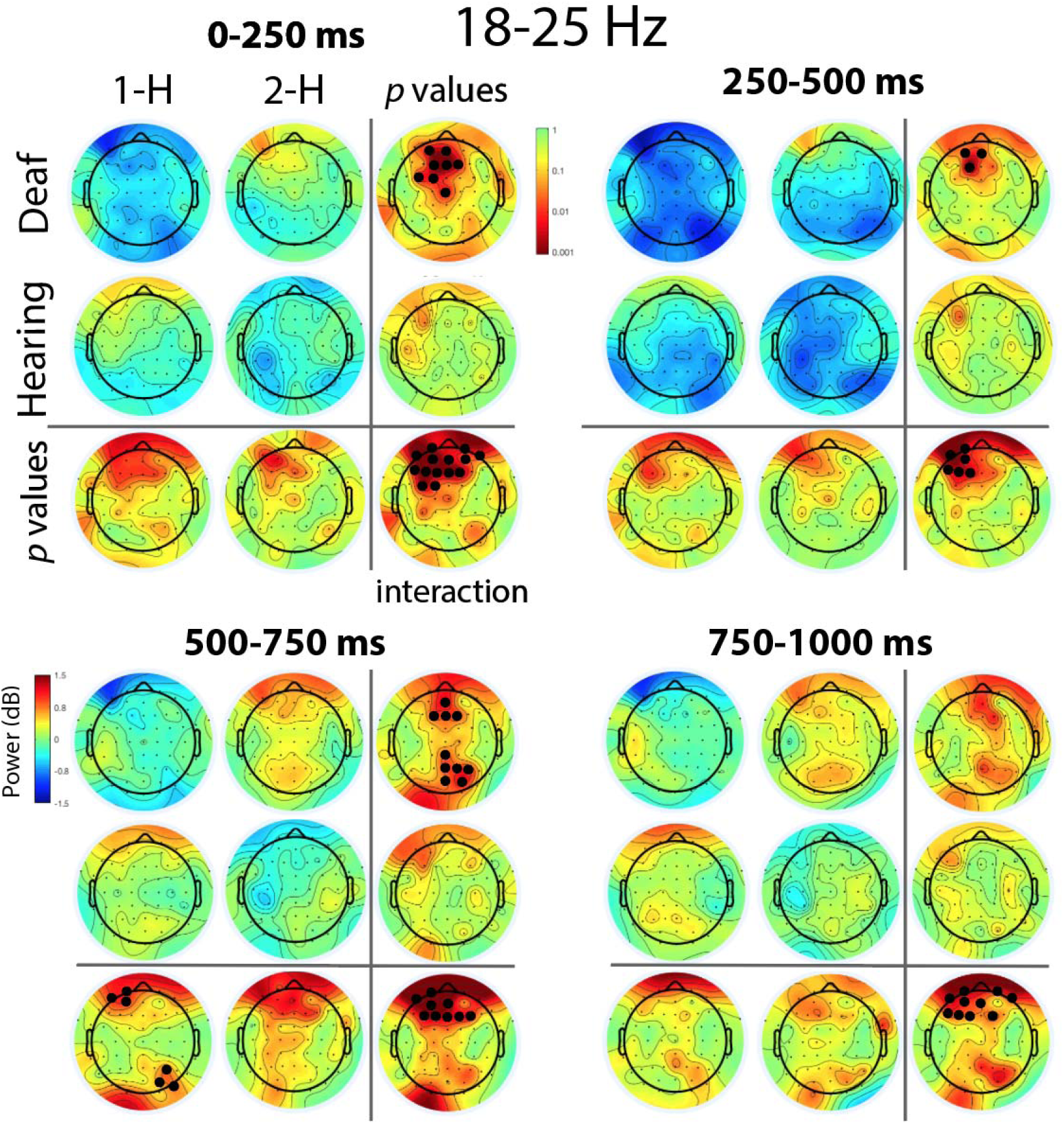
High beta activity (18-25 Hz) across the analysis epoch, 0-1000 ms. Two-way ANOVA comparison (2 Groups: Deaf and Hearing; 2 Conditions: 1-Handed words, 1H and 2-Handed words, 2H) across 64 scalp electrodes, for each of four time bins. Within each time bin, p-values for interaction effects, p-values for main effects, and the average upper-beta power (in dB) for each cell are shown. Electrodes at which the p-value was >.016 are colored in black (minimum cluster size = 3 electrodes). For EEG power maps, cool colors indicate a decrease in power, warm colors indicate and increase in power. For p-value maps, cooler colors indicate high p values, warmer colors indicate lower p-values.

### 3.2 Time-frequency analyses at electrodes in Central region

In the Hearing group, there were no significant effects of Condition at any time in the alpha or beta frequency ranges at any electrode in the central region.

In the Deaf group, two out of 21 electrodes showed significant effects of Condition (1-H vs. 2-H) on alpha power (8-13 Hz) at some point during the analysis epoch (*p <* .05, FDR corrected). For both of these electrodes, there was lower alpha power in the 2H condition compared to the 1-H condition (see Figure 3).

**Figure 3.**
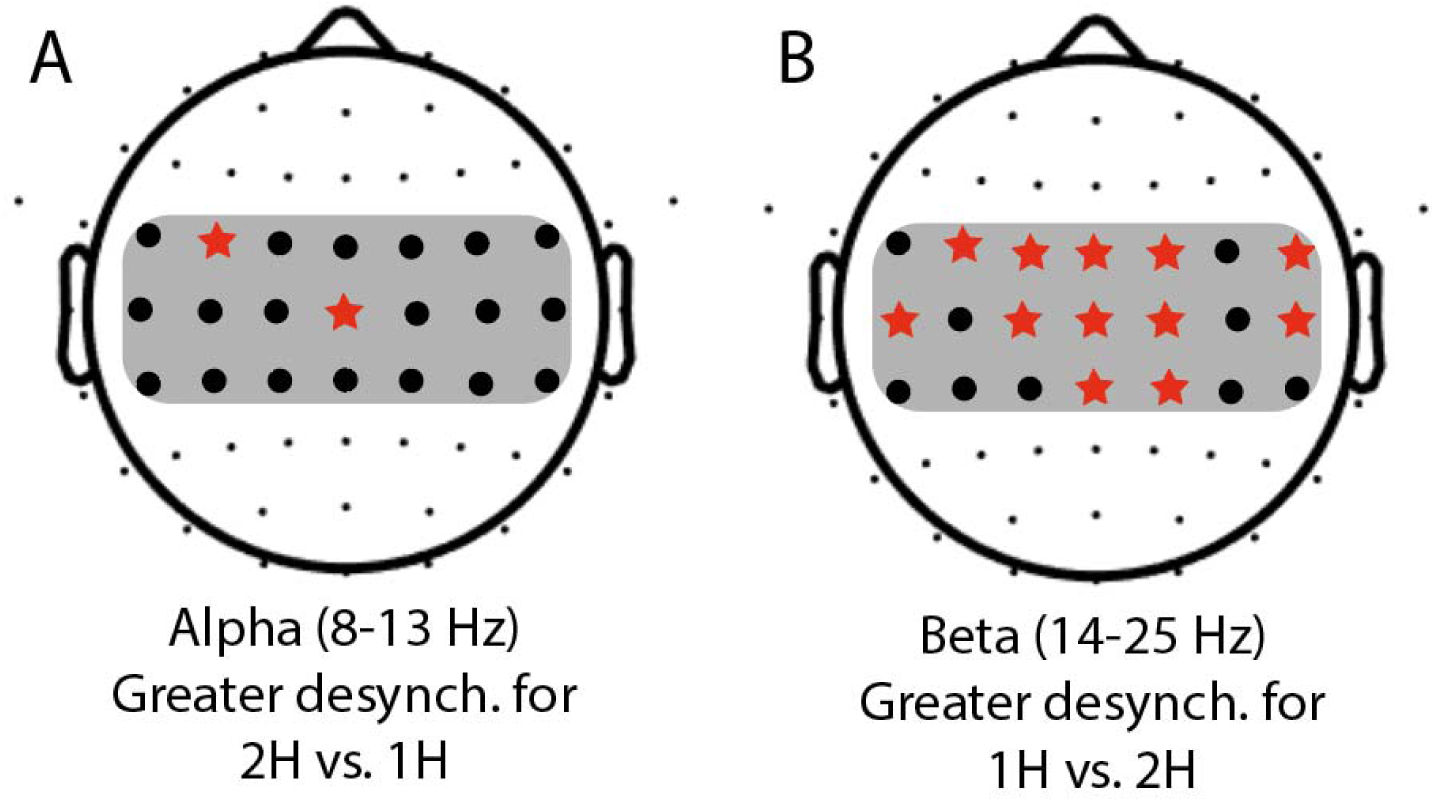
Alpha and beta power across the central region of interest (ROI) in the Deaf group. The 21 electrodes comprising the central ROI are shaded. **A**. For any electrode where alpha (8-13 Hz) power was significantly different during viewing of English words with one- and two-handed (1H and 2H) ASL translations, the electrode is shown with a red star (*p* < .05, FDR corrected). At two electrodes there was significantly greater desynchronization during viewing of 2H words compared to 1H words. B. For any electrode where beta (14-25 Hz) power was significantly different during viewing of English words with 1H and 2H ASL translations, the electrode is shown with a red star (*p* < .05, FDR corrected). At all starred electrodes there was significantly greater desynchronization during viewing of 1H words compared to 2H words.

In the Deaf group, twelve out of 21 central electrodes showed significant effects of Condition (1-H vs. 2-H) on beta power (14-25 Hz) at some point during the analysis epoch (*p <* .05, FDR corrected). For all twelve of these electrodes, the direction of the effect was the same: there was lower beta power in the 1H condition compared to the 2H condition (see Figures 3 and 4).

**Figure 4.**
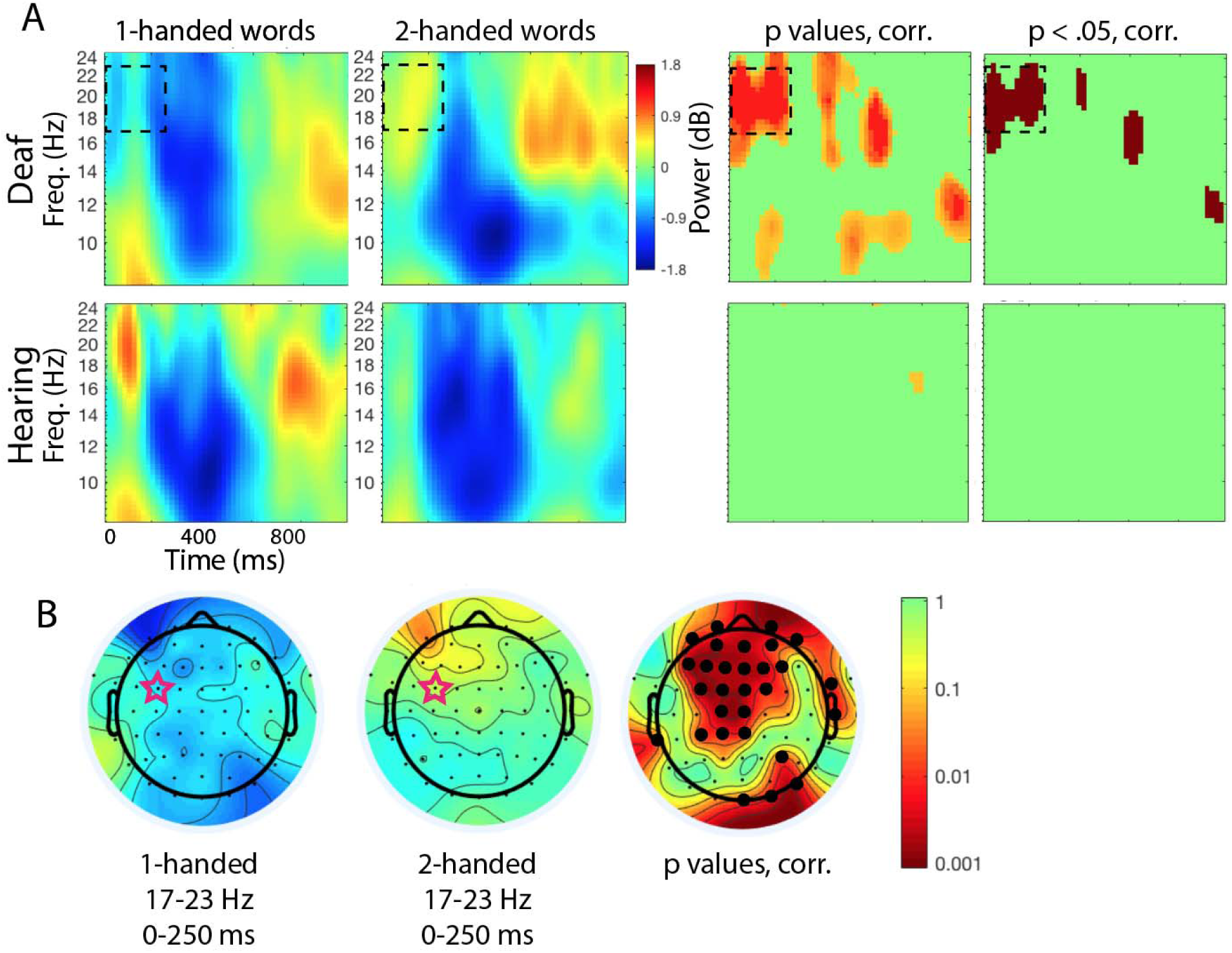
Alpha and beta power during presentation of English words. **A**. Time-frequency representation of ERSPs in response to word onset at time = 0 (duration of word presentation = 1000 ms) for electrode FC3 (starred in Figure 3B). The first two columns show responses to English words with 1- and 2-handed ASL translations, respectively. The third column shows *p* values of the comparison of the first two panels, with the False Discovery Rate (FDR) correction applied. The fourth column shows *p* values thresholded at *p <* .05, FDR corrected. Cool colors indicate a decrease in power, warm colors indicate an increase in power. **B**. Time-frequency responses in the Deaf group across the scalp for 1-handed and 2-handed conditions, at times and frequencies corresponding to the dashed boxes in Figure 3A. Rightmost plot shows the statistical comparison between the conditions; electrodes significant at the *p* < .05 threshold, FDR corrected, are marked with a black circle.

## 4. Discussion

We aimed to investigate if deaf signers simulate sensorimotor properties of ASL signs as they read English words. Based on existing work demonstrating English-to-ASL cross-modal, cross-linguistic translation, we questioned whether neural activity differentiated between English words whose ASL translations had gross sensorimotor differences: how many hands are used to produce the signs. Here, we present evidence that when Deaf signers read, alpha and beta range EEG rhythms respond differently to English words whose ASL translations differ in this way. The pattern of results presented here supports the notion of cross-modal, cross-linguistic translation, and suggests that re-enactment of the actions used to produce the signs constitutes one mechanism involved in this phenomenon.

### 4.1 Central alpha effects

We predicted that there would be significantly greater alpha desynchronization over central electrodes in response to reading 2H words compared to 1H words, based on the greater gross motor effort required to perform a two-handed movement compared to a movement that uses one hand. While our results do support this prediction, the effect was not particularly strong. Two electrodes within the central region showed statistically significant support for this pattern of results. Additionally, several electrodes (e.g., FC3, from 200-800 ms, seen in Figure 4) showed effects that trended toward significance in the alpha rhythm but did not pass our criteria to be considered significant. Nevertheless, the results seen in the central alpha rhythm do provide some evidence that the sensorimotor cortex is more active when a Deaf signer reads English words whose ASL translations use both hands.

It is likely that the relation between number of hands used to produce an action and degree of sensorimotor cortex activity is not entirely straightforward, thus clouding this pattern of results. For instance, the relation between recruitment of sensorimotor cortex and degree of movement may vary depending on the actor’s expertise with the movements (Koeneke, Lutz, Wüstenberg, & Jäncke, 2004; Vogt et al., 2007). One other factor which may affected the direction of our results is frontal inhibition mechanisms reflected by the differences in frontal beta activity between the two conditions. During voluntary production of movements, alpha desynchronization is usually most evident over the contralateral hemisphere, especially in the pre-movement preparation period (Pfurtscheller & Lopes da Silva, 1999). Thus, one might have expected that we would see most evident differences in the alpha rhythm over the right hemisphere, given that the primary difference between the 1H words and the 2H words is the involvement of the left hand (for which the right sensorimotor cortex is contralateral). However, our results did not fit this prediction. This is likely because during action observation (or, in this case, action simulation), the alpha response is bilateral in nature, since there is no pre-movement motor preparation involved (Avanzini et al., 2012). The pre-movement motor preparation during action production is typically a more bilateral response. Overall, the current results do suggest that gross sensorimotor characteristics of ASL signs are activated when deaf signers read English, there is more work to do with regard to comparing various types of sensorimotor characteristics. For instance, future work may more carefully consider the symmetry of ASL sign translations, or the type of movements used when producing the signs.

### 4.2 Fronto-central beta effects

We predicted that reading words with 2-handed ASL translations would result in greater beta desynchronization compared to 1-handed words, due to the typical decrease in beta power when sensorimotor processing demands are increased. However, the experiment yielded significant results going in the opposite direction: reading words with 1-handed translations was associated with greater beta desynchronization. Different effects in different subdivisions of the alpha and beta bands is typical, and perhaps representative of different aspects of stimulus processing (Weiss & Mueller, 2012). While the majority of the sensorimotor beta literature shows desynchronization in situations of greater sensorimotor engagement, plenty of studies have demonstrated the opposite effect of less desynchronization associated with sensorimotor engagement, in line with our current findings (Cheyne et al., 2003; Neuper & Pfurtscheller, 2001; Pfurtscheller, Krausz, & Neuper, 2001; Quandt, Marshall, Bouquet, & Shipley, 2013; Spitzer & Haegens, 2017). In the field as whole, the exact nature of the relationship between beta power and sensorimotor engagement is not entirely clear, but beta rhythms are thought to have an important role in the relationship between sensorimotor processes and language (Spitzer & Haegens, 2017; Weiss & Mueller, 2012).

Our beta EEG results were most evident at frontal and fronto-central electrodes. While both 1H and 2H words resulted in some beta desynchronization, the Deaf group showed significantly less desynchronization when reading English words with a 2-handed ASL translation compared to English words with a 1-handed ASL translation (see Figure 4). And, immediately after stimulus onset, as well as ~600 ms after stimulus onset, a clear beta synchronization (i.e., beta rebound) is evident. Multiple processes may be reflected in these results, including some simulation of the action, as reflected in the beta desynchronization, but also inhibition of covert imitation, seen in the higher beta power for words whose ASL translations would require greater demands on the sensorimotor system. Frontal beta power has been associated with inhibition of motor processes (Alegre et al., 2004; Walsh, Kühn, Brass, Wenke, & Haggard, 2010), suggesting the possibility that the frontal beta effects we report here are due to greater inhibition of motor simulation processing when Deaf signers read 2H English words. The frontal distribution of our beta-range effects may implicate these frontal inhibitory processes related to the blocking of actual movement production during the viewing of 2H words.

An alternate explanation for the frontal beta effects is that mentally simulating one-handed signs actually requires increased sensorimotor control. It may be that although signers show greater primary sensorimotor cortex activity during production of two-handed signs (Emmorey et al., 2016), the processes occurring during mental simulation of signs are quite different. For instance, it is possible that since one-handed signs are inherently asymmetric (one hand is moving and the other is stationary), frontal inhibitory mechanisms are differently recruited for these 1-handed signs as compared to symmetric 2-handed signs. In addition, our results may be influenced by the fact that the 2H word group included both words for which the ASL translation involved a symmetric sign (e.g., FAMILY, with the left and right hands moving symmetrically with identical handshapes) and words for which the ASL translation involved an asymmetrical sign (e.g., KEY, for which both the handshapes and movements differ for each hand). It is likely that action simulation processes vary significantly depending on the symmetry of an imagined action, with symmetrical bimanual actions associated with significantly less sensorimotor cortex activity compared to asymmetrical actions, as has been shown during execution of bimanual actions (Sadato, Yonekura, Waki, Yamada, & Ishii, 1997). This is also likely to be sensitive to the complexity of the actions (Toyokura et al., 2002), presenting yet another factor which should be further studied in future work.

The analysis at the central region of interest revealed significant effects upon the beta rebound effect present in the ERSP. Starting around 600 ms post-stimulus onset, a clear synchronization of beta power is evident in the 2H condition for the Deaf group only (see Figure 4A). This so-called “beta rebound” occurred at most electrodes in the central ROI, always in the same direction with increased beta power in the 2H condition. The beta rebound is a typical characteristic of the sensorimotor beta rhythm, generally following the initial beta desynchronization associated with onset of a performed or observed action (Avanzini et al., 2012; Pfurtscheller & Lopes da Silva, 1999). We had not explicitly predicted a difference in beta rebound activity between the two conditions, but it is interesting to consider its meaning since the effect was quite robust. Beta synchronization and beta rebound are related to onset and stopping of movement (Alegre, Alvarez-Gerriko, Valencia, Iriarte, & Artieda, 2008), and the beta rebound effect is closely linked to activity in the primary motor cortex (Parkes, Bastiaansen, & Norris, 2006). These beta rebound effects, in combination with the alpha-band desynchronization effects seen in response to 2H words, support our original predictions: that reading 2H words would result in greater activity in the primary sensory and motor cortices compared to reading 1H words.

Human and primate work both show that an increase in frontal beta power may also be indicative of maintenance of an item in visual short-term memory, maintenance of a current state of mind, or cognitive control mechanisms more generally (Stoll et al., 2016; Weiss & Mueller, 2012). The robust frontal beta effects we report here may be the result of a number of higher-level cognitive control processes. In addition, we do see evidence for greater activity in primary sensory and motor cortices in response to 2H words, as we originally predicted. Given the basic manipulation of this study, in which we compared EEG responses to English words with ASL translations that varied on large-scale sensorimotor characteristics, we are confident that any inhibitory or control-based mechanisms stem from the somatosensory and motor-related requirements of the hand and body movements required to produce the related ASL signs. The lack of significant differences between the two word groups on English and ASL linguistic parameters lends additional support to this claim. Finally, since no significant effects were observed in the Hearing group, it is clear that these differences reflect a process arising from an individual’s experiences as a Deaf ASL signer.

### 4.3 General discussion

The observed differences in alpha and beta rhythm EEG power occurred in the context of a single-word reading task. Our participants were tasked with keeping track of how many animal words (e.g., bear, duck) they saw, with the intention of ensuring they were processing the semantics of each word. Our findings demonstrate that very soon after word onset, the sensorimotor characteristics (e.g., use of one or two hands) of the ASL translations of these words affected the sensorimotor EEG rhythms—with significant effects within the first 250 ms. Other recent work has shown that the beta rhythm is sensitive to sensorimotor associations with words during a reading task very soon after onset of the stimulus—within 140 ms of word onset (Bechtold, Ghio, Lange, & Bellebaum, 2018), which is consistent with our findings. The current results show that multiple processes affect sensorimotor EEG rhythms when signers read individual English words. This work is the first demonstration of these effects, and future studies in this area could further dissociate between various categories of signs (e.g., symmetrical and asymmetrical two-handed signs) as well as exploring other sensorimotor characteristics of signs beyond the one-hand vs. two-hand distinction. Future studies along these lines can clarify which aspects of action simulation (e.g., motor imagery, tactile mirroring) contribute to cross-modal, cross-linguistic translation in deaf signers.

Future work must also carefully consider the role of ASL and English competence relating to action simulation processes. It has been shown in recent years that signers who are weaker in ASL may rely more heavily on internal representations of the sensorimotor aspects of sign (Corina & Gutierrez, 2016). In our current work, the Deaf group comprised a diverse group of self-identified fluent deaf signers with a range of fluency in ASL and in English (and in some cases, in other written or signed languages).

A next step of this work would restrict participation to participants with more tightly-controlled language backgrounds, in order to statistically assess the role of English fluency and ASL fluency upon the strength of the sensorimotor representations simulated during reading. Our current study did not include a standardized measure of either ASL or English competency, which could provide further objective information to paint a more complete picture of this phenomenon.

## 5. Conclusion

Here we present findings suggesting that sensorimotor characteristics of signs are called upon when Deaf signers read English words. We reveal for the first time the neural oscillatory dynamics occurring during this process, indicating that both alpha and beta EEG oscillations reflect sensorimotor characteristics of ASL signs during cross-modal, cross-linguistic translation. The alpha rhythm showed some desynchronization in response to English words for which the ASL translation uses both hands, compared to words for which the translation uses one hand. This finding was in line with our predictions that such alpha desynchronization would result due to increased activity in the central region overlying the primary sensory and motor cortices. On the other hand, the beta rhythm displayed the opposite pattern, with robust effects showing a pronounced desynchronization in response to “one-handed” English words, compared to “two-handed” words. We suggest this pattern may be due to increased sensorimotor control and inhibition involved in simulating one-handed signs, or due to frontal inhibitory processes related to the blocking of actual movement production during the viewing of 2H words. It is important to keep in mind that the findings in the Deaf group may not be easily interpretable using a literature that largely has drawn from hearing participants; it is possible, and even likely, that neural oscillatory activity during language processing shows notable differences between deaf and hearing populations, much as do other aspects of neurobiology of language (Corina, Lawyer, & Cates, 2012). Our work demonstrates the need for a more complete understanding of how ASL and English overlap and interact with one another in Deaf bilingual readers.

## Acknowledgements

The authors are grateful to Taylor Wardle, Naseem Majrud, Kirsten Daley, and Sawyer Willis for their assistance implementing the study and running participants. We are also thankful for each of the individuals who took the time to participate in this research study.

